# CalciumNetExploreR: An R Package for Network Analysis of Calcium Imaging Data

**DOI:** 10.1101/2025.01.24.634485

**Authors:** Simone Lenci, Dirk Sieger

## Abstract

Analyzing calcium imaging data to understand complex functional networks can be challenging, often requiring multiple tools, custom scripts, and some coding expertise. To address these challenges, we present CalciumNetExploreR (CNER), an R package designed to streamline and standardize the analysis of time-series data from neuronal populations. While originally developed with calcium imaging in mind, CNER’s modular design also makes it adaptable to other sources of neural time-series data, such as fMRI or electrophysiological recordings.

CNER integrates essential steps—normalization, binarization, population activity visualization, network construction, degree distribution analysis, principal component analysis (PCA), power spectral density (PSD) evaluation, and event frequency calculations—into a single, cohesive pipeline. This comprehensive approach enables users to efficiently extract and compare network metrics, including clustering coefficients, global efficiency, community structures, and principal component variances. By offering a flexible and customizable framework, CNER simplifies the examination of functional connectivity and network topology, effectively providing the means to characterize a cellular functional network or analogous structures in other modalities.

Designed as a user-friendly package, CNER allows both experimental and computational neuroscientists to incorporate robust statistical and graphical analyses into their workflows without extensive coding knowledge. By unifying key analytical components into one pipeline, CNER reduces barriers associated with large-scale data analyses, ultimately facilitating deeper insights into the functional organization and dynamic properties of neuronal networks across diverse recording techniques.

## 2 Introduction

Calcium live imaging is a pivotal technique in neuroscience and cellular biology for monitoring intracellular calcium dynamics, which are essential to numerous physiological processes such as neurotransmission, muscle contraction, and signal transduction pathways (Grienberger & Konnerth, 2012; Miyawaki, 2003). By utilizing fluorescent calcium indicators, researchers can visualize and quantify calcium fluctuations within living cells and tissues, providing real-time insights into cellular activity and intercellular communication networks.

The analysis of calcium imaging data poses significant challenges due to the large volume and complexity of the datasets. After calcium traces are extracted from imaging data researchers face the task of interpreting these signals to uncover underlying biological phenomena. This requires sophisticated tools for data preprocessing, statistical analysis, network construction, and visualization to extract meaningful insights.

While several software tools focus on the initial extraction of calcium traces from raw imaging data, such as Suite2p (Pachitariu et al., 2017), CaImAn (Giovannucci et al., 2019), and Fiji (Schindelin et al., 2012)—which can be extended with plugins like TurboReg and Template Matching for motion correction and signal extraction—there is a relative scarcity of tools dedicated to the downstream analysis of these extracted time-series data. Many researchers resort to custom scripts or general-purpose software, which may not be tailored to the specific needs of calcium imaging data analysis. There is a lack of software solutions that enable graph-modeling analyses, such as measuring network clustering, efficiency, and other key topological properties, which are crucial for understanding the dynamic interactions within cellular networks. Existing tools for time-series analysis, such as MATLAB’s Signal Processing Toolbox (The MathWorks, Inc., 2021) and Python libraries like NumPy and SciPy (Harris et al., 2020; Virtanen et al., 2020), offer general-purpose functions but require extensive programming and lack specialized functions. Moreover, they do not provide integrated workflows for network analysis tailored to calcium imaging. Packages like the Brain Connectivity Toolbox (Rubinov et al., 2009) and BRAPH (Mijalkov et al., 2017) offer a wide array of network analysis methods but are not specifically designed for calcium imaging data and often necessitate complex data preparation and manipulation. BRAPH (Brain Analysis using Graph Theory) is a MATLAB-based software package designed for the analysis of brain connectivity using graph theory (Mijalkov et al., 2017). While BRAPH offers extensive functionalities for brain connectivity analysis, it is primarily geared towards processing neuroimaging data such as MRI and fMRI. Applying BRAPH to calcium imaging time-series data may not be straightforward due to differences in data structure and may require significant adaptation or preprocessing steps. Moreover, since BRAPH is built on MATLAB, it requires access to proprietary software, which may not be readily available to all researchers due to licensing costs. This can limit accessibility, especially in educational or resource-constrained settings.

To address these gaps, we introduce CalciumNetExploreR (CNER), an R package designed to streamline the analysis of calcium imaging time-series data with a particular focus on understanding cellular networks and dynamic behaviors in biological systems. CNER is developed in R, an open-source programming language and environment widely used for statistical computing and graphics. By being free to use, it lowers the barrier to entry and promotes wider adoption across the research community. Additionally, R has a vast ecosystem of packages and a strong community support, facilitating collaboration and extension of functionalities.

CNER operates on pre-extracted calcium traces and offers a comprehensive suite of tools for data preprocessing, network analysis, and visualization. CNER offers several key advantages:

- **Preprocessing:** Accepts time-series (calcium fluctuations) and cell/ROI coordinates, with options for normalization and binarization.
- **Network Analysis:** Builds correlation-based networks and extrapolates key features.
- **Subset Analysis:** Isolates user-defined cell groups and measures their interactions with the main network.
- **User-Friendly Workflow:** Features an all-in-one pipeline requiring minimal coding.
- **Modular Design:** Allows for customization and integration with other R packages.

By providing a cohesive platform specifically designed for the downstream analysis of calcium imaging time-series data, CalciumNetExploreR fills a critical niche in the current landscape of analytical tools. Unlike other software that may require extensive programming, lack specialized functions for calcium imaging data, or do not integrate network analysis capabilities tailored to such data, CalciumNetExploreR enables advanced exploratory data analysis with just a few functions. Researchers can isolate a subset of neurons with CalciumNetExploreR to compare their activity and interactions against the larger network, offering insights into specialized functions, cell-type-specific behaviors, and how local changes affect global dynamics. This is especially useful when certain cells are labeled differently (e.g., with a distinct fluorescent reporter) or exposed to targeted stimuli, making it easier to study disease, development, or learning processes.

In this paper, we present the design, functionalities, and applications of CalciumNetExploreR. We highlight its unique capabilities in subset network analysis and the measurement of interactions between main networks and subsets. Through application on sample datasets, we demonstrate how CalciumNetExploreR facilitates advanced analysis and visualization of calcium imaging time-series data, ultimately contributing to a better understanding of complex biological systems.

## 3 Methods

### 3.1. Data Preprocessing

#### 3.1.1 Normalization

To account for potential heterogeneity in baseline levels of the Ca^2+^ indicator (e.g. GCaMP) expression among cells, we normalize the raw calcium signals for each cell individually. This ensures comparability across the cell population by scaling each cell’s signal to a common range and across experiments. The normalization is performed using min-max scaling:

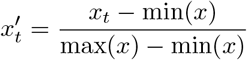

where:

- 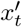 is the normalized calcium signal at time point *t*,
- *x*_*t*_ is the raw calcium signal at time point *t*,
- min(*x*) and max(*x*) are the minimum and maximum calcium signals of the cell over the entire recording period.

This transformation scales the calcium signals of each cell to a range between 0 and 1.

#### 3.1.2 Binarization

The `binarize()` function in *CalciumNetExploreR* is flexible and allows users to define their own criteria for distinguishing between active and inactive states of cells. Users can specify arbitrary thresholds through custom functions to suit their specific experimental conditions. In the context of this study, we binarized the normalized calcium signals based on a threshold derived from the cell’s signal variability. A cell is considered active at time point *t* if its normalized signal exceeds two standard deviations above the mean:

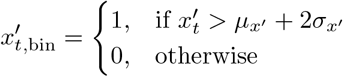

where:

- 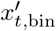 is the binarized signal,
- 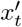 is the normalized signal at time *t*,
- *µ*_*x*_′ is the mean of the normalized signal *x*′,
- *σ*_*x*_′ is the standard deviation of *x*′.

This approach identifies significant calcium events relative to the cell’s baseline activity, allowing for the detection of spontaneous or evoked calcium transients.

### 3.2 Population Activity Analysis

We calculate the proportion of active cells over time by summing the binarized signals across all cells at each time point and dividing by the total number of cells:

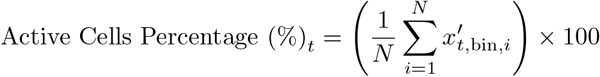

where:

- *N* is the total number of cells,
- 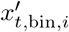 is the binarized signal of cell *i* at time point *t*.

This metric provides insight into the overall network activity and can reveal synchronized activity patterns.

#### 3.2.1 Hierarchical Clustering

To detect potential clusters of calcium activity patterns among cells, we perform hierarchical clustering directly on the normalized calcium signals *x*′ using Ward’s method (Murtagh & Legendre, 2014). This method minimizes the total within-cluster variance and facilitates the identification of groups of cells with similar calcium activity patterns.

### 3.3 Network Creation and Analysis

#### 3.3.1 Functional Connectivity

Functional connectivity between cells is assessed using pairwise cross-correlation of the normalized calcium signals. In practice, `CNER` uses the R function `ccf()` present in the stats package to compute the cross-correlation coefficients. The `ccf()` function calculates the cross-correlation between two time series using the formulae as described in Brockwell and Davis (1991): let 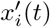 and 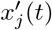 denote the normalized calcium signals of cell *i* and cell *j* at time *t*. The cross-correlation coefficient *ρ*_*ij*_(*τ*) between these two time series at lag *τ* is calculated as:

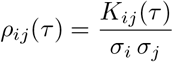

where:

- *K*_*ij*_(*τ*) is the cross-covariance between 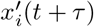 and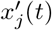;
- *σ*_*i*_ and *σ*_*j*_ are the standard deviations of 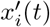 and 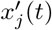, respectively. The cross-covariance *K*_*ij*_(*τ*) is defined as:

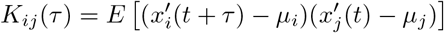

where:

- *µ*_*i*_ and *µ*_*j*_ are the mean values of 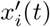 and 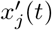, respectively (the time series are modeled as stochastic processes);
- *E* is the expectation value operator. As time in this case is fixed, *E* is the average value from multiple samples.

To account for short delays in cellular responses, cross-correlation coefficients are computed for user-defined lags τ. By default, we consider lags τ = *−*1, 0, +1 time points, but the user can specify different lag values. The maximum absolute value of *ρ*_*ij*_(*τ*) across these lags is selected as the functional connectivity measure between cells:

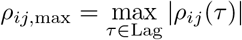

where:

- *ρ*_*ij*_(τ) is the cross-correlation coefficient between cell *i* and cell *j* at lag τ,
- Lag is the set of lags specified by the user.

#### 3.3.2 Network Construction

The functional connectivity matrix *ρ*_*ij*,max_ is used to construct an undirected weighted graph *G* = (*V, E, W*), where:

- *V* is the set of nodes representing cells,
- *E* is the set of edges representing significant functional connections,
- *W* is the set of edge weights given by *ρ*_*ij*,max_.

A threshold *θ* is applied to *ρ*_*ij*,max_ to determine significant edges:

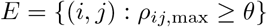

Edges with weights below the threshold are excluded to focus on the most significant connections.

#### 3.3.3 Graph Metrics

We compute several key topological properties of the network to characterize its structure:

- **Degree** (*k*_*i*_): The degree of node i is the number of edges connected to it:

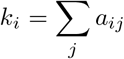

where *a*_*ij*_ = 1 if there is an edge between nodes *i* and *j*, and 0 otherwise.

- **Clustering Coefficient** (*C*(*g*)): Measures the tendency of the entire network to form tightly knit clusters. It is defined as the average clustering coefficient across all nodes in the network:

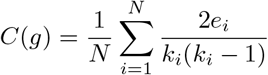

where *e*_*i*_ is the number of edges between the neighbors of node *i, k*_*i*_ is the degree of node *i*, and *N* is the total number of nodes in the network.

- **Global Efficiency (***G*(*g*)): Reflects how efficiently information is exchanged over the entire network. It is calculated as:

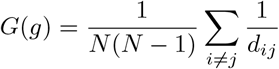

where *d*_*ij*_ is the shortest path length between nodes *i* and *j*, and *N* is the total number of nodes in the network.

- **Mean Degree**: The average degree across all nodes:

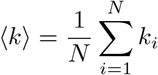

- **Community Detection**: Communities within the network are identified using the leading eigenvector method (Newman, 2006), which calculates the leading non-negative eigenvector of the modularity matrix of the graph. This method partitions the network into clusters of densely connected nodes by maximizing the modularity.

These metrics provide insights into the network’s topology, revealing patterns of connectivity and potential functional modules within the cellular network.

### 3.4 Subset Network Analysis

The package allows for the selection and analysis of subsets of cells within the global network. Subsets can be defined based on various criteria, such as anatomical location, functional characteristics, or genetic markers.

For each subset, a functional network is constructed using the same methods described above. The topological properties of the subset network are calculated and compared to those of the main network. Interactions between the subset and the main network are assessed by analyzing the connections between cells in the subset and those outside it.

This analysis provides valuable information about the role of specific cell groups within the overall network and how they influence or are influenced by the broader cellular community.

### 3.5 Principal Component Analysis (PCA)

*CNER* performs PCA on the normalized calcium signals using the `prcomp` function from the `stats` R package. This process reduces dimensionality and identifies principal components that capture the most variance in the data.

Scree plots are generated to visualize the variance explained by each principal component, aiding in the selection of components for further analysis.

### 3.6 Power Spectral Density (PSD) Analysis

We conduct PSD analysis to examine the frequency content of the calcium signals, identifying dominant frequencies and oscillatory patterns. We use the Welch method (Welch, 1967) to estimate the PSD for each cell’s normalized signal.

The PSD plots display power as a function of frequency, allowing for the identification of periodic behaviors and synchronization phenomena within the cellular population.

### 3.7 Integrated Analysis Pipeline

An integrated analysis pipeline is provided through the `pipeline()` function in the CalciumNetExploreR package. This function automates the sequential execution of the analysis steps with minimal user intervention. The pipeline includes:

1. **Data Loading**: Reads the time-series data and cell coordinates provided by the user.
2. **Normalization and Binarization**: Applies the normalization and binarization procedures as described above.
3. **Population Activity Visualization**: Generates plots of the proportion of active cells over time, with options for hierarchical clustering to detect activity patterns.
4. **Network Construction and Visualization**: Builds the functional network using cross-correlation coefficients and visualizes it using graph plotting functions. Node positions are determined by the provided cell coordinates.
5. **Graph Metrics Calculation**: Computes key topological properties of the network, including degree distribution, clustering coefficient, global efficiency, and identifies communities using the leading eigenvector method.
6. **Subset Analysis**: Allows for the selection of cell subsets for specialized analysis, repeating the network construction and metric calculations for the subset.
7. **PCA and PSD Analyses**: Performs PCA and PSD analyses on the normalized signals, providing visualizations such as scree plots and PSD graphs.
8. **Reporting**: Compiles the results into summary tables and generates comprehensive visualizations for interpretation.

The pipeline function enhances reproducibility and efficiency by encapsulating the analysis workflow into a single command, which can be customized with function arguments to suit specific research needs.

### 3.8 Software and Implementation

The *CalciumNetExploreR* package is developed using R (version 4.4.0) and can be found, with its documentation, on https://github.com/simo-91/CalNetExploreR.

### 3.9 Data Requirements

The package requires:

- **Time-Series Data**: A matrix or data frame where rows represent cells and columns represent time points, containing the raw or pre-processed calcium signals. Mathematically, this can be represented as *M*_*n*×*t*_, where *n* is the number of cells (rows) and *t* is the number of time points (columns). Each element *M*_*ij*_ represents the calcium signal of cell *i* at time point *j*. It is essential that this matrix be properly formatted so that each row corresponds to a unique cell and each column to a time point in the recording.
- **Cell Coordinates**: A matrix or data frame containing the spatial coordinates of each cell (e.g., X and Y positions), necessary for network visualization and spatial analyses. This is represented as a matrix *C*_*n*×2_, where *n* is the number of cells and each row contains the (*x*_*i*_, *y*_*i*_) coordinates of cell *i*. Proper formatting ensures that each cell’s coordinates are aligned with the rows of the time-series matrix, so that spatial information corresponds directly to the calcium data.
- **Labels**: An additional column in the cell coordinates data frame that specifies the subset classification of each cell. This column, named `Label`, contains identifiers (a categorical label) indicating whether a cell belongs to a specific subset.

### 3.10 Experimental Design

#### 3.10.1 Zebrafish Maintenance

Zebrafish (*Danio rerio*) were maintained at 28.5°C on a 14-hour light/10-hour dark cycle in the aquatics facility at BVS Aquatics department based at the Queen Medical Research Institute (University of Edinburgh). Embryos were obtained by natural spawning from adult transgenic *Tg(NBT:H2B-GCaMP6s; nacre*^*-/-*^) and raised in E3 embryo medium at 28.5°C. To inhibit pigmentation in pigmented strains, embryos were treated with 200*µ*M 1-phenyl-2-thiourea (PTU) (Sigma) starting at 24 hours post-fertilization until the end of the experiment. All animal procedures were approved by the Institutional Animal Care and Use Committee (IACUC) and conducted in accordance with the Animal (Scientific Procedures) Act 1986 and institutional guidelines.

#### 3.10.2 Zebrafish oocyte injections and sample generation

We utilized a transgenic zebrafish line *Tg(NBT:H2B-GCaMP6s; nacre*^*-/-*^), in which all neurons express the nuclear calcium indicator GCaMP6s, enabling the visualization of neuronal activity in vivo. To achieve mosaic overexpression of the oncogene in the brain, embryos at the single-cell stage were coinjected with a *pDEST-NBT:*Δ*lexPR-lexOP-pA* driver plasmid and either a *Tol2-pDEST-lexOP:AKT1-lexOP:tagRFP* plasmid for the AKT1 condition (TREAT) or a *Tol2-pDEST-lexOP:tagRFP-pA* control plasmid for the control condition (CTRL). Injection solutions contained 40 ng/uL of plasmid DNA, 0.2% Phenol Red dye and Tol2 capped mRNA (20 ng/µL) to enable transient mosaic expression. Briefly, the transcriptional activator ΔlexPR is expressed in neurons and activates gene downstream of lexOP sites. Hence, this approach resulted in a subset of neurons overexpressing AKT1 and being labeled with red fluorescent protein (RFP) for the TREAT condition and neurons simply being labeled with tagRFP for the CTRL condition.

#### 3.10.3 Calcium Live Imaging

Six days post-fertilization zebrafish larvae were immobilized by treatment with 0.5 mg/mL Mivacurium Chloride (Abcam, ab143667) to inhibit movement. The larvae were then embedded in 1.5% low melting point agarose (Life Technologies) within glass-bottom dishes (MatTek) filled with E3 medium. Timelapse calcium imaging was performed using a Zeiss LSM880 confocal microscope equipped with a 20x water-dipping objective lens, focusing on the hindbrain area. Time-lapses lasted for 15 minutes, capturing frames every 2 seconds (resulting in a 0.5 Hz acquisition rate). Fluorescence excitation was achieved using 488 nm laser lines for GCaMP6s and 543 nm for tagRFP.

### 3.11 Statistical Analysis

Statistical analyses are conducted using built-in R functions and validated methods. When comparing network metrics between groups or conditions, appropriate statistical tests (e.g., t-tests, ANOVA) are applied, and p-values are adjusted for multiple comparisons where necessary.

## 4 Results

### 4.1 Explorative and comparative analysis of network features using CNER

To demonstrate the capabilities of *CalciumNetExploreR* (*CNER*), we explored Ca^2+^ imaging data from exemplary datasets generated using larval zebrafish. We utilized a transgenic zebrafish line *NBT:H2B-GCaMP6s; nacre*^*-/-*^, in which all neurons express the nuclear calcium indicator GCaMP6s, enabling the visualization of neuronal activity in vivo. To perform a comparative analysis using CNER, we generated two experimental datasets: 1) A control dataset (CTRL) in which in addition to the GCaMP6s signal a subset of neurons expressed RFP, and a treatment dataset (TREAT) in which a subset of neurons expressed RFP and the human oncogene AKT1 (see *Methods* for details) (Chia et al., 2018, 2019). As AKT1 overexpression has been shown to influence Ca^2+^ levels before, we deemed this dataset suitable for a comparative analysis using CNER. Datasets consisted of calcium fluorescence time-series data from hundreds of neurons recorded over T = 450 time points (Fig. 1). Finally, we applied the pipeline() function integrated in CNER to the control (CTRL) and the treatment (TREAT) datasets in order to compare their network features. The features analyzed included:

**Figure 1:**
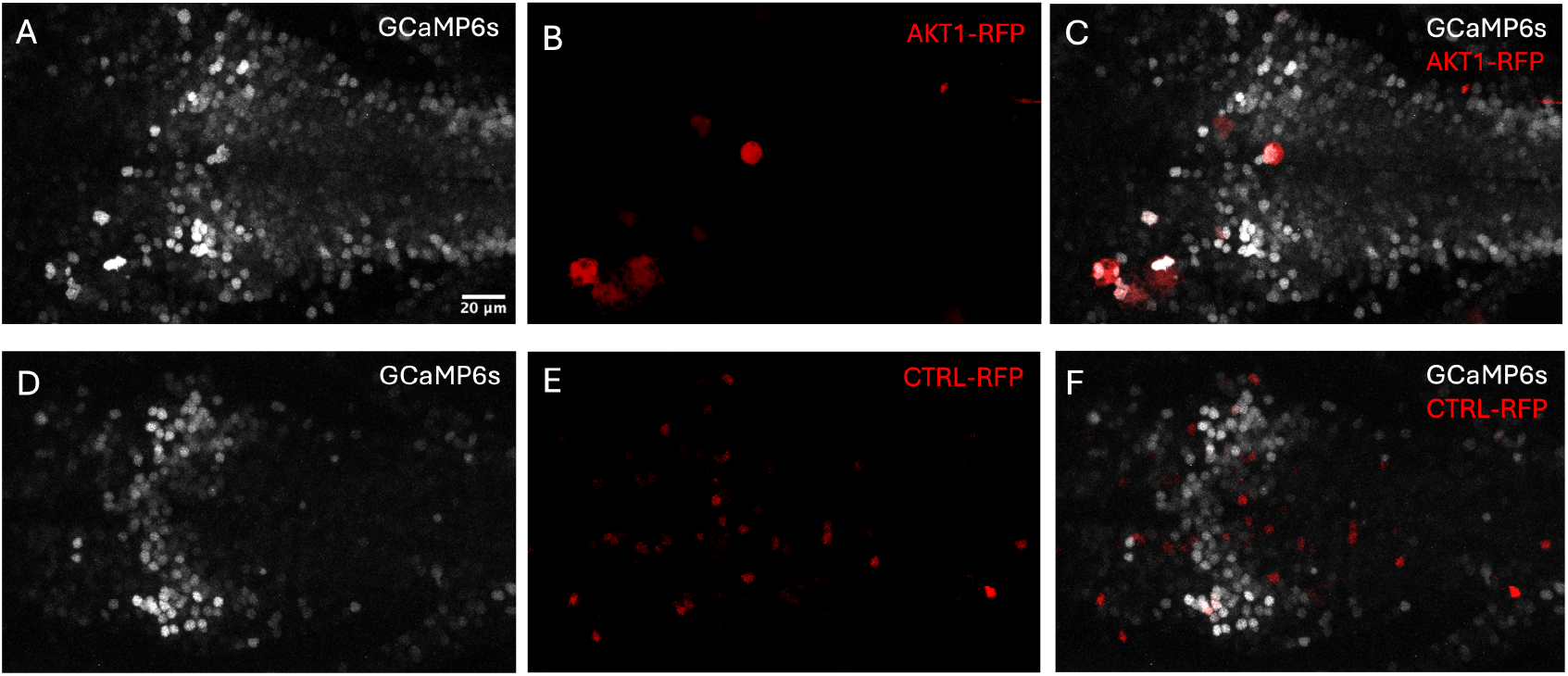
Two-channel calcium imaging time-lapses from the hindbrain of a 6 dpf larval zebrafish in treatment (TREAT, A–C) and control (CTRL, D–F) conditions. The zebrafish line *Tg(NBT:H2B-GCaMP6s;nacre*^*-/-*^) expresses nuclear GCaMP6s in all neurons (grey, GCaMP6s-GFP channel), enabling visualization of neuronal activity. Mosaic expression of either an AKT1 oncogene construct or a control construct was achieved by coinjecting *pDEST-NBT:*Δ*lexPR-lexOP-pA* at the singlecell stage along with either *Tol2-pDEST-lexOP:AKT1-lexOP:tagRFP* (TREAT, AKT1-RFP^+^ neurons in red, panels B,C) or *Tol2-pDEST-lexOP:tagRFP-pA* (CTRL, neurons carrying control plasmid in red, panels E,F). (A,D) GCaMP6s-GFP channel; (B,E) RFP channel; (C,F) merged channels.

- Mean event frequency of calcium events per minute for the entire population (`Freq.`)
- Mean event frequency per minute for the labeled subset of neurons (`Freq. labeled`), which are the neurons overexpressing AKT1 or carrying the control plasmid.
- Clustering coefficient (*C*_*g*_)
- Global efficiency (*G*_*g*_)
- Percentage of variance explained by the top five principal components (`Top5PC Var (%)`)
- Proportion of connections from labeled-to-unlabeled neurons over total possible labeled-to-unlabeled connections (`LtU`)

The `pipeline()` function is a comprehensive analysis tool that automates the processing of calcium imaging data through several key stages, integrating all of the following functions that allow for customization and detailed analysis. It essentially acts as a wrapper for several functions present in the package, namely `normalize()`, `binarize()`, `population_activity()`, `make_network()`, `plot_network()`, `degrees()`, `pca()`, `PSD.plt()`, `get_top5pc_variance()`, `subset_connections()`, `events_per_min()` and also igraph functions `transitivity()` and `global_efficiency()`.

The results are summarized in Table 1.

**Table 1:** Network Features for AKT1 and CTRL Samples.

| Sample | Condition | Top5PC Var (%) | $C_g$ | $G_g$ | LtU | Freq. | Freq. labeled |
| --- | --- | --- | --- | --- | --- | --- | --- |
| Sample 1 | CTRL | 12.14 | 0.775 | 0.01394 | 0.012072 | 2.886 | 2.92 |
| Sample 2 | CTRL | 13.50 | 0.760 | 0.01500 | 0.013500 | 3.000 | 2.96 |
| Sample 3 | CTRL | 11.50 | 0.780 | 0.01250 | 0.010632 | 2.750 | 2.78 |
| Sample 4 | TREAT | 8.91 | 0.547 | 0.00931 | 0.006000 | 2.209 | 4.00 |
| Sample 5 | TREAT | 9.10 | 0.536 | 0.01000 | 0.005600 | 2.150 | 4.10 |
| Sample 6 | TREAT | 8.40 | 0.557 | 0.00850 | 0.006400 | 2.250 | 3.90 |

Firstly, the datasets were normalized using the `normalize()` function, integrated in the pipeline, which scales the raw calcium signals across cells to ensure comparability by centering and scaling to unit variance.

Next, we converted the normalized, continuous signals into binary states (active/inactive) using the `binarize()` function to facilitate the analysis of neuronal activity patterns.

#### 4.1.1 Pipeline() function unravels latent features and differences between treatment and control networks

One of the outputs of the `pipeline()` function is the mean event frequency per minute for the entire population of neurons (`Freq.`). According to the analyzed samples, CTRL samples had a Freq. ranging between 2.75 and 3.00 events/min, while TREAT samples showed lower values (around 2.15–2.25 events/min). This difference was statistically significant (*p* <0.01, Fig.2A). Another output of the pipeline is the mean event frequency per minute for the labeled subset of neurons (Freq. labeled). The TREAT samples showed a significantly higher mean event frequency (4.00 ± 0.10) compared to CTRL samples (2.89 ± 0.09), representing an increase of approximately 39% (*p* < 0.001, independent t-test, Fig. 2B). This suggests that the treatment enhances neuronal excitability or firing rate, which is reflected in the increased frequency of calcium events detected.

**Figure 2:**
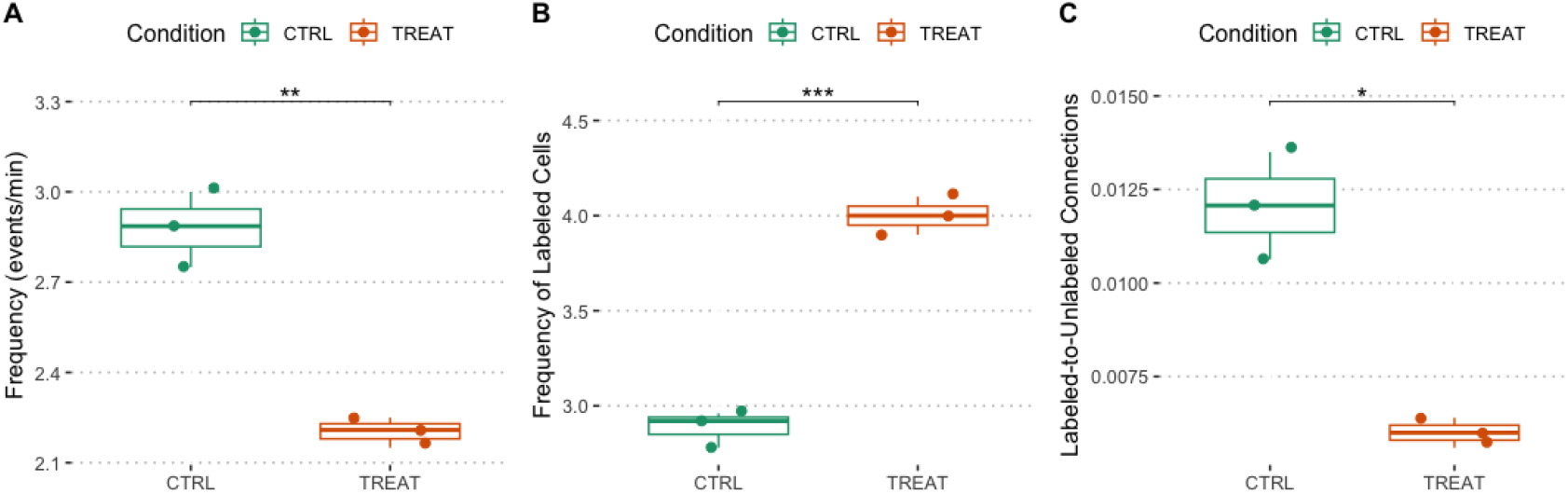
CNER revealed differences in neuronal activity and connectivity patterns between treatment and control conditions. (A) Frequency of events (`Freq.`) for the entire population of neurons in control (CTRL) and treatment (TREAT) conditions. TREAT samples exhibited significantly lower Freq. values compared to CTRL samples (Freq_CTRL_ = 2.75–3.00 events/min; Freq_TREAT_ = 2.15–2.25 events/min; *p* < 0.01, independent t-test). (B) Frequency of labeled neurons (Freq_labeled_) between control (CTRL) and treatment (TREAT) conditions. A significant increase in Freq_labeled_ was observed in the TREAT condition compared to CTRL (Freq_CTRL,labeled_ = 2.89 ± 0.09; Freq_TREAT,labeled_ = 4.00 ± 0.10; *p* < 0.001, independent t-test). (C) Proportion of connections from labeled to unlabeled neurons (LtU) in CTRL and TREAT conditions. A significant decrease in LtU was observed in the TREAT condition compared to CTRL (LtU_CTRL_ = 0.01207 ± 0.00143; LtU_TREAT_ = 0.00600 ± 0.00040; *p* < 0.05, Mann-Whitney U test).

The `subset_connections()` function integrated within the `pipeline()` quantified the proportion of connections between specified subsets of neurons. We found that the proportion of connections from labeled to unlabeled neurons (`LtU`) was higher in CTRL samples (0.01207 ± 0.00143) compared to TREAT samples (0.00600 ± 0.00040), indicating a two-fold decrease in the TREAT condition (*p* < 0.05, MannWhitney U test, Fig. 2C). This result suggests that neurons in the treatment condition are less connected to the surrounding neuronal network, potentially leading to functional isolation or altered network integration.

To construct functional connectivity graphs, the pipeline uses the `make_network()` function, which performs cross-correlation analysis on the binarized calcium signals to infer connections between neurons. Customization options in this function allowed us to adjust parameters such as `lag.max` (numeric, default 1) specifying the maximum lag for cross-correlation, and `correlation_threshold` (numeric or `“none”`, default 0.3) to filter edges by weight; setting it to `“none”` disables filtering. This flexibility enabled us to tailor the sensitivity of the network construction based on correlation strength and lag. For this use case, we used a correlation threshold of 0.6 and a lag of 1.

Visualization of the networks was done using the integrated `plot_network()` function, which displays the network structure and highlights community memberships by labeling nodes accordingly. Customization options included coordinates (a data frame with columns `“X”`, `“Y”`, `“Cell”`, and `“Label”)`, `cell_- ID` (character, options `“cell”` or `“communities”)` to label nodes by cell ID or community, and `label` (logical, default `FALSE`) to include labels on the plot.

In the TREAT condition, the network exhibited reduced clustering and global efficiency compared to the CTRL condition. The mean clustering coefficient (*C*_*g*_) in TREAT samples was 0.547±0.011, significantly lower than in CTRL samples (0.772 ± 0.010), representing a 41% decrease (*p* = 0.0009, independent t-test, Fig. 3A). Global efficiency (*G*_*g*_), calculated using built-in functions from the igraph package, was reduced by 33% in TREAT samples (0.00927 ± 0.000750) compared to CTRL samples (0.0138 ± 0.00125) (*p* = 0.004, independent t-test, Fig. 3B). These metrics indicate that the treatment leads to a less interconnected and less efficient network.

**Figure 3:**
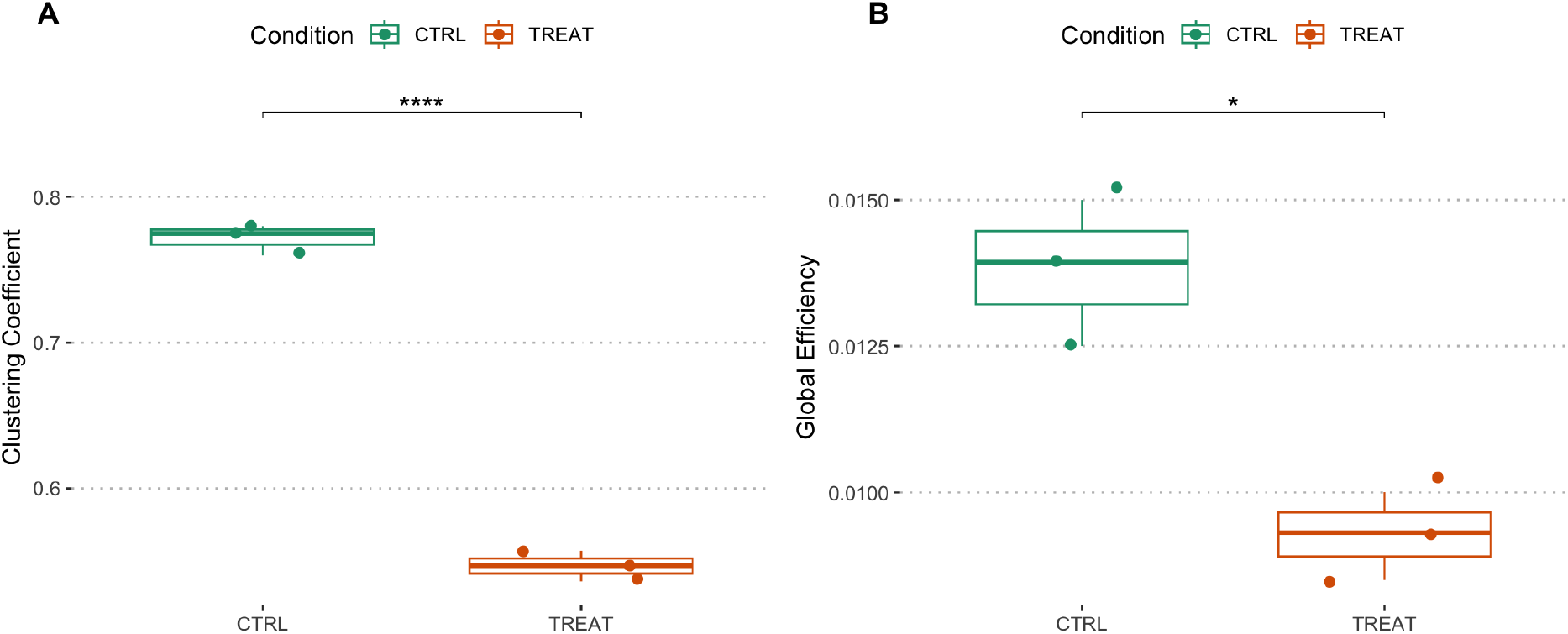
Quantification of Clustering Coefficient and Global Efficiency in networks using CNER. Treatment reduces clustering and network efficiency in the zebrafish brain. (A) Clustering coefficient (*C*_*g*_) and (B) Global efficiency (*G*_*g*_) between control (CTRL) and treatment (TREAT) conditions. Significant reductions in both *C*_*g*_ and *G*_*g*_ were observed in the TREAT condition compared to CTRL (*C*_*g*CTRL_ = 0.772 ± 0.010; *C*_*g*TERAT_ = 0.547 ± 0.011; *p* = 0.0009. *G*_*g*CTRAL_ = 0.0138 ± 0.00125; G_*g*TREAL_ = 0.00927 ± 0.000750; *p* = 0.004, independent t-test).

The network graphs generated by the `plot_network()` function, integrated in the pipeline, visually demonstrate these differences (Fig. 4). The TREAT network (Fig. 4A) shows reduced clustering with only one large community (defined as a community with more than five members), whereas the CTRL network (Fig. 4B) displays increased clustering with 14 large communities. Nodes are color- and number-coded by community membership, illustrating the fragmentation in the TREAT condition.

**Figure 4:**
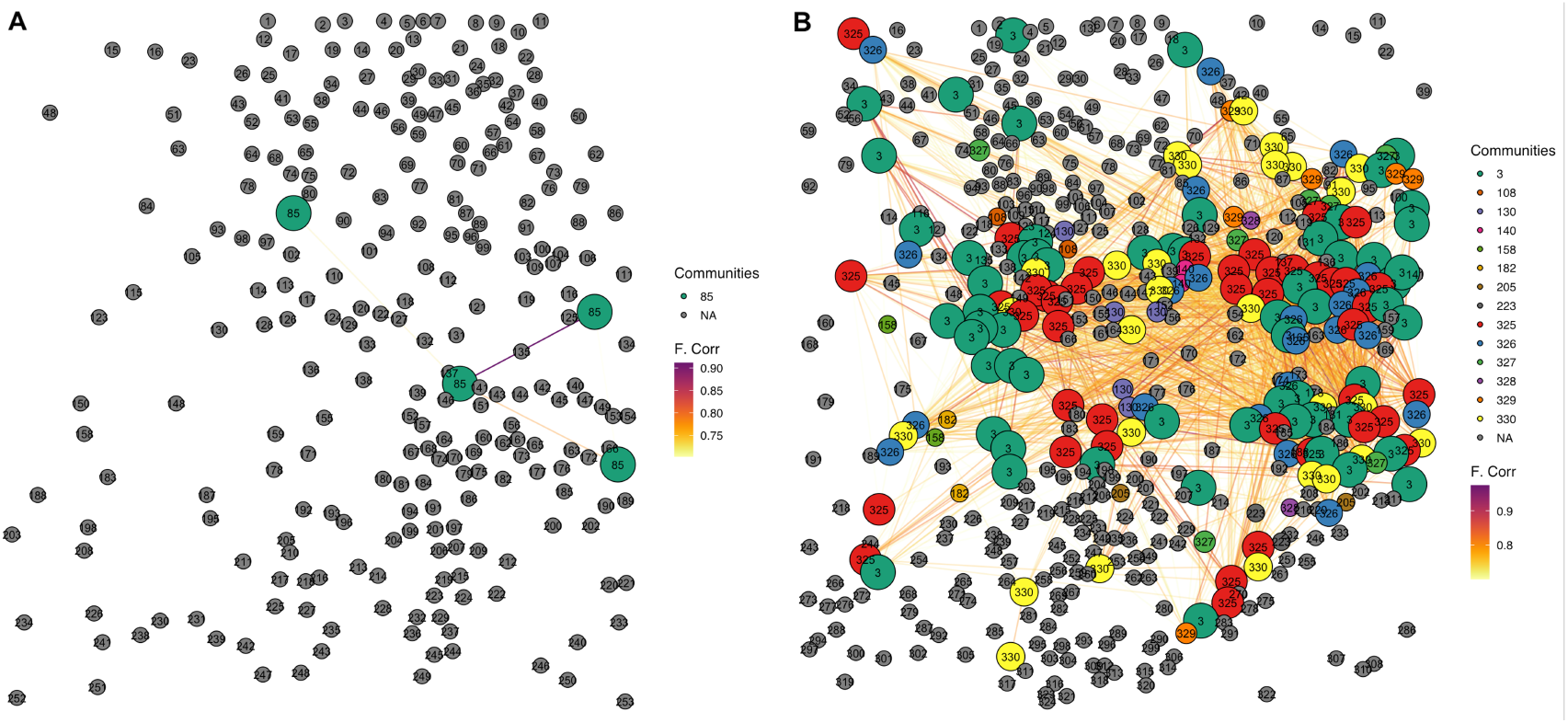
Network graphs generated reveal reduced clustering and lower number of big communities in the treatment condition. Comparison of network graphs for treatment (TREAT, A) and control (CTRL, B) conditions generated by the `plot_network()` function. The TREAT network exhibits reduced clustering with only one large community, while the CTRL network displays increased clustering with 14 large communities. Nodes are color- and number-coded by community membership.

We visualized population activity using the population_activity.plt() function, which generates raster plots of neuronal activity over time, sorted through hierarchical clustering to reveal patterns of co-activation. Customization options included the dendrogram argument (logical, default FALSE) to include a dendrogram in the plot. In the TREAT condition, the plots showed increased variability and less organized activity patterns compared to CTRL (Fig. 5), suggesting a lack of functional clusters and disrupted synchronization among neurons in the treatment group.

**Figure 5:**
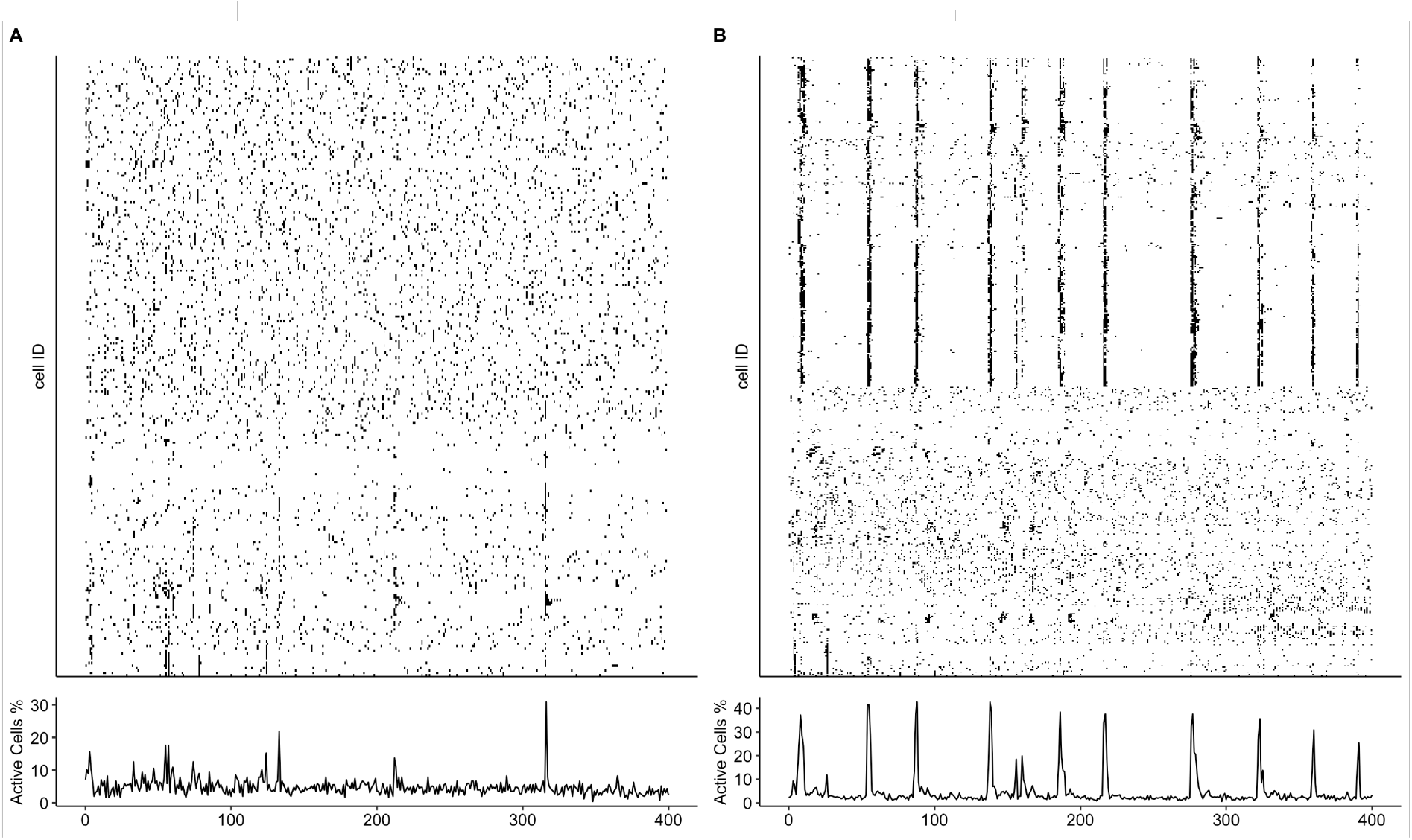
Population activity plots generated by CNER reveal lack of functional clusters in the treatment condition. Population activity in representative samples for TREAT (A) and CTRL (B) conditions, visualized using the `population_activity.plt()` function. The TREAT condition shows increased variability and less organized activity patterns compared to CTRL.

We assessed network complexity using the `pca()` function also integrated within the `pipeline()`.

This function performs principal component analysis to identify the principal components that capture the most variance in the data, thereby reducing dimensionality and highlighting significant patterns of neuronal activity. The percentage of variance explained by the top five principal components (`Top5PC Var (%)`) was significantly lower in TREAT samples (8.80% ± 0.35%) compared to CTRL samples (12.38% ± 1.02%), indicating a reduction of approximately 29% (*p* < 0.01, independent t-test). This suggests that neuronal activity in the TREAT condition is more complex and less dominated by a few principal patterns.

The PCA scree plots generated by the `pca()` function illustrate this difference in network complexity (Fig. 6). In the TREAT condition (Fig. 6A), the variance explained by each successive principal component declines more gradually, indicating that a larger number of components are needed to capture the same amount of variance as in the CTRL condition (Fig. 6B).

**Figure 6:**
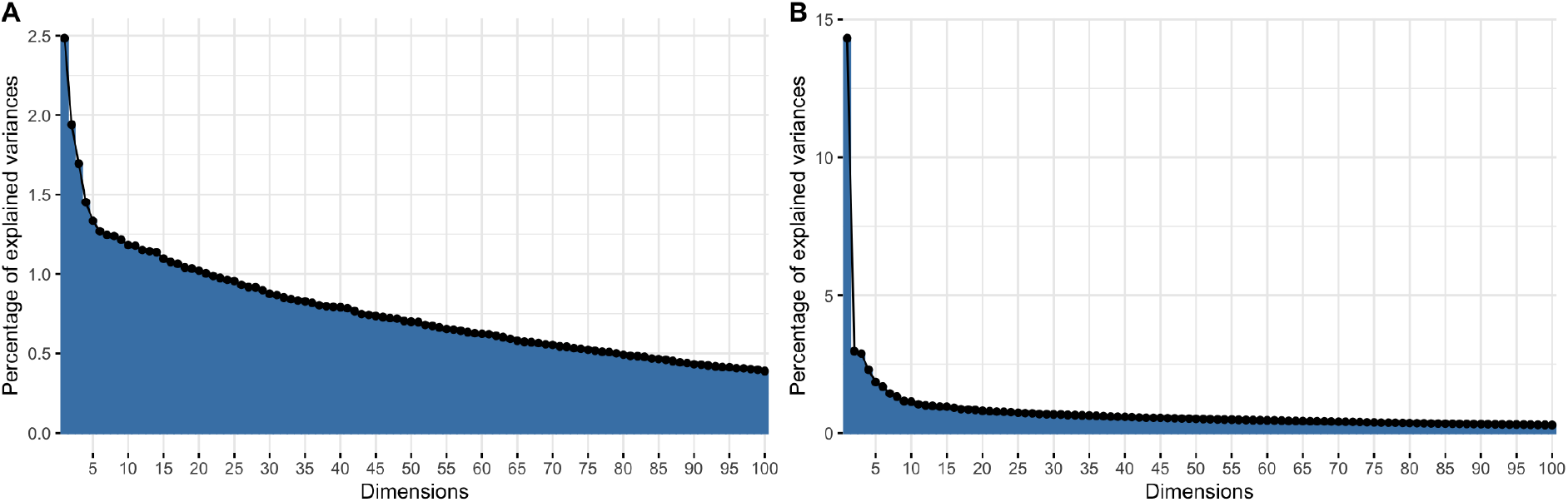
PCA analysis shows increases network complexity in treatment group. Representative scree plots after PCA for TREAT (A) and CTRL (B) conditions, generated using the `pca()` function. The CTRL network is explained by fewer principal components, suggesting lower complexity than the TREAT network, which retains higher variance across more components (Percentage of variance explained by the top five principal components; CTRL, mean: 12.38% ± 1.02%; TREAT, mean: 8.80% ± 0.35%; *p* < 0.01, independent t-test).

By utilizing the `pipeline()` wrapper function of *CalciumNetExploreR*, we effectively analyzed and visualized differences in neuronal network features between the TREAT and CTRL conditions. Significant alterations associated with the treatment were identified, demonstrating the utility of *CalciumNetExploreR* in uncovering functional network changes in calcium imaging data.

## 5 Discussion

In this study, we introduced *CalciumNetExploreR* (*CNER*), a comprehensive R package designed for the analysis of calcium imaging data and the exploration of neuronal network features. By applying *CNER* to calcium imaging datasets from zebrafish larvae under AKT1 overexpression (TREAT) and control (CTRL) conditions, we demonstrated the package’s capability to extract meaningful network metrics and visualize complex neuronal interactions. The use of the `pipeline()` function allowed for a simple and standardized analysis workflow, encompassing data normalization, binarization, network construction, feature extraction, and visualization.

By analyzing the TREAT and CTRL datasets, we were able to characterize significant differences in network topology and function using *CNER*. Specifically, the package revealed that in the TREAT condition there is a substantial decrease in the clustering coefficient (*C*_*g*_) and global efficiency (*G*_*g*_). The mean *C*_*g*_ was reduced by approximately 41% in the TREAT condition compared to CTRL, indicating a disruption in local connectivity and network clustering. Similarly, the global efficiency decreased by about 33%, suggesting impaired overall network integration and efficiency in information transfer. The network graphs generated by *CNER* (Fig. 4) visually illustrated the reduced clustering and altered community structures in the TREAT networks compared to CTRL. Additionally, the population activity plots provided insights into the temporal dynamics of neuronal activity, revealing increased noise and variability in the TREAT condition networks (Fig. 5). The application of principal component analysis (PCA) within the package offered insights into the complexity of neuronal activity. By analyzing the variance explained by principal components, we found that the top five principal components accounted for a lower percentage of variance in the TREAT condition compared to CTRL. This suggests increased complexity and heterogeneity in neuronal firing patterns in the treatment group, which could be further investigated using *CNER*’s tools. Furthermore, *CNER* facilitated the analysis of subpopulations within the neuronal network. By examining the connections from labeled (RFP^+^) to unlabeled neurons and calculating the mean event frequencies, we observed that neurons of the treatment condition had elevated activity levels and altered connectivity patterns. Specifically, the mean event frequency per minute for labeled neurons was significantly higher in the TREAT condition, and the proportion of connections from labeled to unlabeled neurons was decreased. These analyses demonstrate how *CNER* can be used to explore the functional roles of specific neuronal subpopulations within the broader network. Importantly, these findings highlight how *CNER* can uncover alterations in latent network properties that may be associated with experimental manipulations or disease models.

Graph theory, on which *CNER* is based, has emerged as a powerful framework for modeling and analyzing cellular population activity in neuroscience, as it provides quantitative tools to characterize complex neuronal networks and interpret functional connectivity patterns derived from calcium imaging data (Blevins et al., 2022; Bullmore & Sporns, 2009; Nelson & Bonner, 2021; Rubinov & Sporns, 2010; Sporns et al., 2004). Several elegant studies have demonstrated the effectiveness of graph theoretical approaches in elucidating network structures and dynamics (Avitan et al., 2017; Malmersjo et al., 2013; Smedler et al., 2014; Tibau et al., 2013; Wei et al., 2020).

Finally, the adaptability and extensibility of *CNER* allow users to incorporate additional analyses or customize functions to suit their specific research needs. The package’s reliance on widely used R packages such as `igraph`, `ggplot2`, and ggraph ensures compatibility and ease of integration with custom analytical workflows.

In summary, our application of *CalciumNetExploreR* to the TREAT and CTRL datasets demonstrates how the package can be utilized to characterize and compare neuronal networks under different experimental conditions. By providing comprehensive tools for network analysis and visualization, *CNER* enables researchers to uncover subtle changes in network topology and function, contributing valuable insights into the mechanisms underlying neuronal dynamics.

### 5.1 Limitations and Future Directions

Despite its capabilities, *CalciumNetExploreR* has limitations that warrant consideration. The package currently assumes a basic level of proficiency in R programming and familiarity with network analysis concepts, which may present a barrier for some users. A graphical user interface (GUI) is in the works, as it could improve accessibility for a broader range of researchers.

Additionally, while the package offers several network metrics and visualization options, the incorporation of a baseline correction in pre-processing and advanced machine learning tools could further enrich the analyses. Future developments may focus on expanding the package’s functionality to include these features, thereby providing a more comprehensive tool for neuroscientific research.

## 6 Conclusion

*CalciumNetExploreR* is a powerful and versatile R package that facilitates the analysis of calcium imaging data and the exploration of neuronal network properties. By streamlining complex analytical processes and providing intuitive visualization tools, it enables researchers to uncover meaningful insights into neuronal dynamics and network architecture. Our application of *CalciumNetExploreR* to AKT1 overexpression data illustrates its utility in identifying significant differences in network features between experimental conditions. We believe that *CalciumNetExploreR* will be a valuable resource for the neuroscience community, promoting advanced network analyses and contributing to a deeper understanding of neuronal function.

## Acknowledgements

The authors thank the Bioresearch and Veterinary Services (BVS) Aquatics facility (QMRI, University of Edinburgh) for maintenance and care of the zebrafish. The authors are grateful to Dr. Daniel Soong and the UK Zebrafish Imaging and Screening Facility for assistance with microscope imaging. Special thanks to Dr. Nicola Roman’o for critical reading of this article.

## Ethics

Animal experiments were reviewed and approved by the ethical review committee of the University of Edinburgh and conducted under Home Office project license PP8119722, in accordance with the Animals (Scientific Procedures) Act 1986.

## Key points

- CalciumNetExploreR (CNER) is an R package designed for straight-forward, user-friendly analysis and visualization of calcium live imaging data, providing a powerful tool for studying neuronal networks in neuroscience.
- CNER integrates preprocessing, network analysis, graph modeling, and visualization tools, such as population activity plots, functional connectivity networks, degree distributions, and PCA scree plots, into a cohesive pipeline tailored for experimental calcium imaging data.
- CNER enables easy characterization of functional networks and allows the analysis and comparison of subsets within the general neuronal population.
- CNER is flexible and user-friendly, making it accessible to experimental and computational neuroscientists for analyzing calcium imaging data or other time-series datasets.

## References

Avitan, L., Pujic, Z., Mölter, J., Van De Poll, M., Sun, B., Teng, H., Amor, R., Scott, E. K., & Goodhill, G. J. (2017). Spontaneous Activity in the Zebrafish Tectum Reorganizes over Development and Is Influenced by Visual Experience. Current biology: CB, 27 (16), 2407–2419.e4. 10.1016/j.cub.2017.06.056

Blevins, A. S., Bassett, D. S., Scott, E. K., & Vanwalleghem, G. C. (2022). From calcium imaging to graph topology. Network Neuroscience, 6 (4), 1125–1147. 10.1162/netna00262

Brockwell, P. J., & Davis, R. A. (1991, February 22). Time Series: Theory and Methods: Theory and Methods. Springer Science & Business Media.

Bullmore, E., & Sporns, O. (2009). Complex brain networks: Graph theoretical analysis of structural and functional systems. Nature Reviews Neuroscience, 10 (3), 186–198. 10.1038/nrn2575

Chia, K., Keatinge, M., Mazzolini, J., & Sieger, D. (2019). Brain tumours repurpose endogenous neuron to microglia signalling mechanisms to promote their own proliferation (R.M. White, T. Taniguchi, & J.-P. Levraud, Eds.). eLife, 8, e46912. 10.7554/eLife.46912

Chia, K., Mazzolini, J., Mione, M., & Sieger, D. (2018). Tumor initiating cells induce Cxcr4-mediated infiltration of pro-tumoral macrophages into the brain (R. M. White, Ed.). eLife, 7, e31918. 10.7554/eLife.31918

Giovannucci, A., Friedrich, J., Gunn, P., Kalfon, J., Brown, B. L., Koay, S. A., Taxidis, J., Najafi, F., Gauthier, J. L., Zhou, P., Khakh, B. S., Tank, D. W., Chklovskii, D. B., & Pnevmatikakis, E. A. (2019). CaImAn: An open source tool for scalable Calcium Imaging data Analysis. eLife, 8, e38173. 10.7554/eLife.38173

Grienberger, C., & Konnerth, A. (2012). Imaging Calcium in Neurons. Neuron, 73 (5), 862–885. 10.1016/j.neuron.2012.02.011

Harris, C. R., Millman, K. J., van der Walt, S. J., Gommers, R., Virtanen, P., Cournapeau, D., Wieser, E., Taylor, J., Berg, S., Smith, N. J., Kern, R., Picus, M., Hoyer, S., van Kerkwijk, M. H., Brett, M., Haldane, A., del Río, J. F., Wiebe, M., Peterson, P., … Oliphant, T. E. (2020). Array programming with NumPy. Nature, 585 (7825), 357–362. 10.1038/s41586-020-2649-2

Malmersjo, S., Rebellato, P., Smedler, E., Planert, H., Kanatani, S., Liste, I., Nanou, E., Sunner, H., Abdelhady, S., Zhang, S., Andang, M., El Manira, A., Silberberg, G., Arenas, E., & Uhlen, P. (2013). Neural progenitors organize in small-world networks to promote cell proliferation. Proceedings of the National Academy of Sciences, 110 (16), E1524–E1532. 10.1073/pnas.1220179110

Mijalkov, M., Kakaei, E., Pereira, J. B., Westman, E., & Volpe, G. (2017). BRAPH: A graph theory software for the analysis of brain connectivity. PLoS ONE, 12 (8), e0178798. 10.1371/journal.pone.0178798

Miyawaki, A. (2003). Visualization of the spatial and temporal dynamics of intracellular signaling. Developmental Cell, 4 (3), 295–305. 10.1016/s1534-5807(03)00060-1

Murtagh, F., & Legendre, P. (2014). Ward’s Hierarchical Agglomerative Clustering Method: Which Algorithms Implement Ward’s Criterion? Journal of Classification, 31 (3), 274–295. 10.1007/s00357-014-9161-z

Nelson, C. J., & Bonner, S. (2021). Neuronal Graphs: A Graph Theory Primer for Microscopic, Functional Networks of Neurons Recorded by Calcium Imaging. Frontiers in Neural Circuits, 15. 10.3389/fncir.2021.662882

Newman, M. E. J. (2006). Finding community structure in networks using the eigenvectors of matrices. Physical Review E: Statistical Physics, Plasmas, Fluids, and Related Interdisciplinary Topics, 74 (3), 036104. 10.1103/PhysRevE.74.036104

Pachitariu, M., Stringer, C., Dipoppa, M., Schröder, S., Rossi, L. F., Dalgleish, H., Carandini, M., & Harris, K. D. (2017, July 20). Suite2p: Beyond 10,000 neurons with standard two-photon microscopy. 10.1101/061507

Rubinov, M., Kötter, R., Hagmann, P., & Sporns, O. (2009). Brain connectivity toolbox: A collection of complex network measurements and brain connectivity datasets. NeuroImage, 47, S169. 10.1016/S1053-8119(09)71822-1

Rubinov, M., & Sporns, O. (2010). Complex network measures of brain connectivity: Uses and interpretations. NeuroImage, 52 (3), 1059–1069. 10.1016/j.neuroimage.2009.10.003

Schindelin, J., Arganda-Carreras, I., Frise, E., Kaynig, V., Longair, M., Pietzsch, T., Preibisch, S., Rueden, C., Saalfeld, S., Schmid, B., Tinevez, J.-Y., White, D. J., Hartenstein, V., Eliceiri, K., Tomancak, P., & Cardona, A. (2012). Fiji: An open-source platform for biological-image analysis. Nature Methods, 9 (7), 676–682. 10.1038/nmeth.2019

Smedler, E., Malmersjö, S., & Uhlén, P. (2014). Network analysis of time-lapse microscopy recordings. Frontiers in Neural Circuits, 8, 1–10. 10.3389/fncir.2014.00111

Sporns, O., Chialvo, D. R., Kaiser, M., & Hilgetag, C. C. (2004). Organization, development and function of complex brain networks. Trends in Cognitive Sciences, 8 (9), 418–425. 10.1016/j.tics.2004.07.008

The MathWorks, Inc. (2021). Signal processing toolbox user’s guide. manual. Natick, Massachusetts. https://www.mathworks.com/products/signal.html

Tibau, E., Valencia, M., & Soriano, J. (2013). Identification of neuronal network properties from the spectral analysis of calcium imaging signals in neuronal cultures. Frontiers in Neural Circuits, 7. 10.3389/fncir.2013.00199

Virtanen, P., Gommers, R., Oliphant, T. E., Haberland, M., Reddy, T., Cournapeau, D., Burovski, E., Peterson, P., Weckesser, W., Bright, J., van der Walt, S. J., Brett, M., Wilson, J., Millman, K. J., Mayorov, N., Nelson, A. R. J., Jones, E., Kern, R., Larson, E., … van Mulbregt, P. (2020). SciPy 1.0: Fundamental algorithms for scientific computing in Python. Nature Methods, 17 (3), 261–272. 10.1038/s41592-019-0686-2

Wei, Z., Lin, B.-J., Chen, T.-W., Daie, K., Svoboda, K., & Druckmann, S. (2020). A comparison of neuronal population dynamics measured with calcium imaging and electrophysiology. PLOS Computational Biology, 16 (9), e1008198. 10.1371/journal.pcbi.1008198

Welch, P. (1967). The use of fast Fourier transform for the estimation of power spectra: A method based on time averaging over short, modified periodograms. IEEE Transactions on Audio and Electroacoustics, 15 (2), 70–73. 10.1109/TAU.1967.1161901

